# Phenome-wide burden of copy number variation in UK Biobank

**DOI:** 10.1101/545996

**Authors:** Matthew Aguirre, Manuel Rivas, James Priest

## Abstract

Copy number variations (CNV) represent a significant proportion of the genetic differences between individuals and many CNVs associate causally with syndromic disease and clinical outcomes. Here, we characterize the landscape of copy number variation and their phenome-wide effects in a sample of 472,228 array-genotyped individuals from the UK Biobank. In addition to population-level selection effects against genic loci conferring high-mortality, we describe genetic burden from syndromic and previously uncharacterized CNV loci across nearly 2,000 quantitative and dichotomous traits, with separate analyses for common and rare classes of variation. Specifically, we highlight the effects of CNVs at two well-known syndromic loci *16p11.2* and *22q11.2*, as well as novel associations at *9p23*, in the context of acute coronary artery disease and high body mass index. Our data constitute a deeply contextualized portrait of population-wide burden of copy number variation, as well as a series of known and novel dosage-mediated genic associations across the medical phenome.

## Introduction

Copy number variants (CNV) are a class of structural variation often defined as large deletions or duplications of at least 1 kilobase (kb) of genomic sequence. CNVs exhibit substantial variability in both size and frequency in the population and have been implicated across a variety of common and rare health outcomes^1^. Regional deletion and duplication syndromes have also been described at many loci, clustering near microsatellite repeats or areas of segmental duplication which may potentiate non-allelic homologous recombination^2^. For example, CNV-based architectures for neuropsychiatric (e.g. autism spectrum disorder), developmental (e.g. *16p11.2*)^3,4^, and syndromic cardiac disease (e.g. *22q11.2*)^5^ phenotypes have been well established.

Despite a growing body research on CNV-related syndromes and disease etiologies, large-scale studies of CNV effects have been limited by their rarity in the general population. However, burden testing methods which address this rarity by pooling observed variation across gene regions have yielded reproducible associations in the context of congenital heart disease and various neurocognitive outcomes^6,7^. Moreover, as studies which include either microarray or NGS-based genotype data have grown in size and scope, it has become possible to describe the distribution of CNVs at kilobase-level resolution in the general population^8,9^. Furthermore, the aggregation of richly annotated phenotype data in biobanks has diversified the set of phenotypes available for well-powered association studies, and allows for more precise characterization of syndromic CNV-associated disease^10,11,12^.

We here describe the landscape of copy number variation and their associations with 1,937 phenotypes in a cohort of 472,228 participants from the UK Biobank^13^. We replicate well-established syndromic effects of common CNVs — namely *22q11.2* deletion (DiGeorge) syndrome and two variants of *16p11.2* deletion syndrome — and highlight known and novel associations for body mass index (BMI), acute coronary artery disease (CAD), and related adipose and cardiovascular phenotypes. Summary statistics from traditional genome-wide associations for common CNVs, as well as from gene-level aggregate burden tests of rare variants across all phenotypes are available for download on the Global Biobank Engine^14^.

## Results

### Landscape of common and rare CNVs in a large volunteer cohort

To call copy number variants in UK Biobank, we apply PennCNV^15^ separately within each genotyping batch, resulting in 278,455 unique CNVs among 472,228 individuals after sample quality control. We also observe heavy-tailed distributions in size and allele count of CNVs, with average CNV length ~226*kb* and the majority of called variants singleton in the cohort (Figure 1a,b). This translates to notable burden of variation for nearly all individuals, with 439,464 (93.1%) of the individuals possessing at least one CNV detectable at kilobase-resolution (Figure 1c,1d). Among individuals with at least one CNV, we estimate an average burden of 5.5 variants covering >200*kb* of genomic sequence (median 3 variants affecting ~100*kb*, Figure 1c,d). While in-line with previous reports^8^, these estimates of individual-level burden are likely conservative, as regions where array markers are sparse or missing limit the accuracy of variant calling. Furthermore, we are unable to call smaller (<1*kb*) variants due to inconsistent marker density across all chromosomal regions on the Axiom and BiLEVE UK Biobank genotyping arrays. This limitation is visible in the histogram of called CNV lengths (Figure 1a); we call substantially fewer variants on the order of hundreds of base-pairs than on the order of thousands.

**Figure 1:**
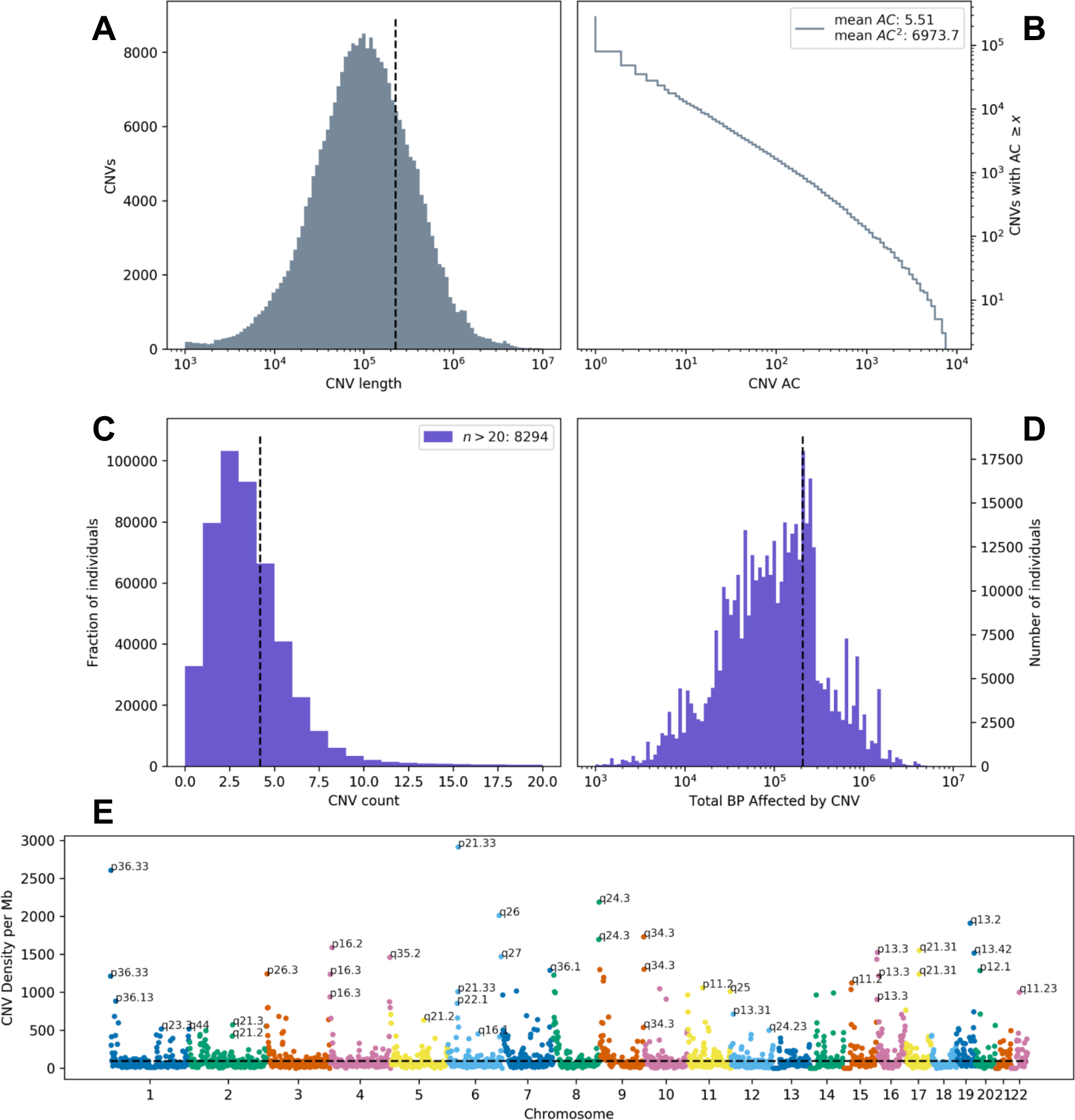
Burden and distribution of copy number variation in UK Biobank. **(A)** Log-scale histogram of CNV lengths. Mean length (dashed line) is 226.5*kb*. **(B)** Cumulative density of CNV allele count (AC), displayed in log-log axes. Average AC is 5.5, but average frequency as experienced by the population (weighted by count, hence AC^2^) is ~1.6%. **(C)** Histogram of CNV counts and **(D)** log-scale base-pairs affected by CNV per individual. Sample-level burden is heavy-tailed, with the average individual carrying 4.2 variants (dashed line), affecting mean ~207.6kb of genomic sequence. **(E)** Genome-wide density of CNV, defined as the number of unique CNVs overlapping 10 megabase (Mb) windows tiling each chromosome. Hotspots of structural variation are labeled by cytogenic band.

We also observe a highly non-uniform burden of variation across genomic position, with breakpoints most common near the ends of chromosomes, and at known regions of segmental duplication (Figure 1e). Among them are *1p36*, *8q24.3*, *9q34.3*, and *19q13*, all of which have associated microdeletion syndromes causing developmental delay with uniquely characteristic growth patterns^16–19^. Other CNV-hotspots like *6p21.33*, which contains the major histocompatibility complex protein gene family, may be influenced by high marker density (in this case for HLA allelotyping) in addition to these biological features which underlie structural mutagenesis. However, these loci do not categorically correspond to areas where structural variation is commonly observed in the population (Figure S1). For example, *1p36* and *19q13* are also the respective sites of common CNVs overlapping *RHD* and *FUT2* (Rhesus and Lewis blood groups), but there are no such common variants within the telomeric *16p13* cytoband.

### Survivorship bias due to genetic selection against early-onset diseases

We estimate gene-level intolerance to structural variation by adapting a method for estimating regional selective constraint^8^. Relative to the general population, the volunteers within the UK Biobank are described to have a “healthy-cohort” enrollment bias^20^ and were enrolled between the ages of 40 to 69, which informs our findings. Within the tail of positive constraint *z-*scores, which indicate the strongest intolerance to structural variation, we observe enrichment for genes which cause early onset diseases, particularly cancer. Among the top fifteen constrained genes (Table 1) are *BRCA1* and *BRCA2*, which are associated with early-onset breast cancer^21,22^; *MLH1*, *MSH2*, *MSH6*, which cause early onset colorectal cancer (Lynch syndrome)^23–25^; and *ATM* and *APC*, which are involved with mismatch repair cancers^26,27^.

**Table 1:** **(Left)**15 genes most intolerant to copy number variation. Columns are gene label, constraint *z*-score, and probability of CNV intolerance (pLI, see Methods for definitions).

|  | Constraint z | CNV pLI |
| --- | --- | --- |
| BRCA2 | 3.262 | 0.9905 |
| BRCA1 | 2.470 | 0.9830 |
| APC | 1.997 | 0.9421 |
| ATM | 1.196 | 0.9886 |
| MSH2 | 1.192 | 0.9875 |
| MLH1 | 1.176 | 0.9959 |
| MSH6 | 0.918 | 0.9928 |
| SBDS | 0.875 | 0.9739 |
| RB1 | 0.872 | 0.9552 |
| SPATA31D1 | 0.869 | 0.9979 |
| CYP3A4 | 0.863 | 0.9979 |
| OTOP1 | 0.848 | 0.9930 |
| PABPC3 | 0.847 | 0.9923 |
| KRT16 | 0.844 | 0.9979 |
| ZNF302 | 0.844 | 0.9979 |

Selections from the most highly constrained pathways from Gene Ontology Consortium^28^ resources (Table 2) also suggest strong intolerance to CNV for genes involved with core biological processes like protein binding, cellular structural integrity (keratinization), development (growth hormone receptor binding), and immune regulation (natural killer cell activation). Similar results at the gene- and pathway-level are observed for deletion-specific constraint (Table S1,S2), whereas duplication-specific analysis suggests autoimmune-related genes and pathways are most strongly intolerant to dosage effects (Table S1,S2). These results indicate strong selective effects occurring prior to enrollment in the UK Biobank during childhood and early adulthood against loss of function variation in core developmental, metabolic, and tumor-suppressing genes, and against dosage-altering variation in immune-related genes.

**Table 2:** **(Below):**15 pathways most enriched for constrained genes (t-test, gene set members versus all others). Columns are GO pathway ID/name, change in *z*-score between set and non-set members, indicating mean strength of selective effect in the pathway, and *p*-value.

| | Pathway | $\Delta z$ | P |
| --- | --- | --- | --- |
| GO:0000137 | Golgi cis cisterna | 0.42 | $1.16 \times 10^{-31}$ |
| GO:0045095 | keratin filament | 0.26 | $7.29 \times 10^{-31}$ |
| GO:0052697 | xenobiotic glucuronidation | 0.49 | $1.03 \times 10^{-21}$ |
| GO:0005515 | protein binding | 0.07 | $1.24 \times 10^{-21}$ |
| GO:0031424 | keratinization | 0.18 | $1.51 \times 10^{-21}$ |
| GO:0000800 | lateral element | 0.46 | $3.46 \times 10^{-21}$ |
| GO:0008194 | UDP-glycosyltransferase activity | 0.36 | $6.13 \times 10^{-21}$ |
| GO:0015020 | glucuronosyltransferase activity | 0.35 | $1.32 \times 10^{-17}$ |
| GO:0008202 | steroid metabolic process | 0.28 | $9.31 \times 10^{-17}$ |
| GO:0005132 | type I interferon receptor binding | 0.35 | $1.78 \times 10^{-16}$ |
| GO:0008274 | gamma-tubulin ring complex | 0.39 | $2.66 \times 10^{-16}$ |
| GO:0002323 | natural killer cell activation involved in immune response | 0.36 | $1.54 \times 10^{-14}$ |
| GO:0005131 | growth hormone receptor binding | 0.54 | $4.33 \times 10^{-14}$ |
| GO:0052696 | flavonoid glucuronidation | 0.45 | $5.40 \times 10^{-14}$ |
| GO:0042954 | lipoprotein transporter activity | 0.40 | $5.78 \times 10^{-14}$ |

### Association testing identifies CNVs at novel and syndromic loci

We compute genome-wide associations across 1,893 phenotypes for 8,274 common CNVs observed at 0.005% allele frequency (1 in 20,000) in our cohort, using regression as implemented in the analysis software PLINK^29^. We also perform L1-regularized regression for rare-variant burden tests, pooled by gene. For these tests, we measure the net effect of rare CNVs (AF < 0.1%) overlapping within 10*kb* of the gene region as defined by HGNC^30^ for 7,614 protein coding genes with at least 5 individuals affected by such a variant. A complete list of phenotypes analyzed is available in Table S3. Here, we describe representative results for one common disease and one quantitative measure with established genetic risk factors and large sample sizes in UK Biobank: acute coronary artery disease (CAD) and body mass index (BMI).

For Acute CAD, we identify one statistically significant (p < 6×10^−6^) association after Bonferroni correction for the common CNV GWAS: an intergenic deletion at chromosome *9p23*. Intergenic variants at the *9p21* locus have been implicated in previous association studies of blood-based biomarkers relevant to cardiac outcomes, specifically, decreases in hematocrit and hemoglobin concentration^31^, as well as carotid plaque burden^32^. A recent meta-analysis^33^ using data from UK Biobank and CARDIoGRAMplusC4D identified a lead variant in the vicinity of this locus (rs2891168) associated with 6% unit increase in risk for similarly defined coronary artery disease. However, the CNV we here identify confers an estimated 12.4-fold increased risk (95%CI: 4.3-35.9, *p*=3.6×10^−6^) and is at least 2Mb distant from the nearest SNPs (rs10961206) at genome-wide significance near the *9p21*/*9p23* locus in the meta-analysis. This and the absence of linkage between the 9p23 CNVs and rs10961206 (*r* = 0.013) are suggestive of independent effects.

Gene-level burden testing of rare CNVs in individuals with CAD implicates *LDLRAD3*, a member of the low density lipoprotein (LDL) receptor family. CNVs called in this gene are predominantly deletions affecting the coding sequence — in aggregate (*n=27*), these variants confer an estimated 10-fold increase in risk of Acute CAD (95% CI: 3.9-25.6, *p*=1.4×10^−6^). Though the role of lipoprotein receptors in cholesterol metabolism is a well established mechanism of risk for cardiovascular disease, *LDLRAD3* is not known to participate in cholesterol metabolism. It is, however, a receptor widely expressed throughout adult tissues which may participate in proteolysis in the central nervous system^34,35^. We therefore sought to replicate these findings using two-sample mendelian randomization^36^ on expression quantitative trait loci (eQTLs) from CAD summary statistics from a CARDIoGRAMplusC4D meta-analysis^37^. We identify a nominally significant protective effect between an eQTL increasing expression of *LDLRAD3* and CAD (OR=0.85 [95%CI: 0.62-0.97], *p*=0.012), the direction of which is consistent with a dosage model of *LDLRAD3*-mediated risk for CAD.

We also find two significant associations for BMI, both deletions at chromosome *16p11.2*, a locus implicated in syndromic early onset obesity and developmental delay. Each of these CNVs appears to correspond to a distinct form of *16p11.2* deletion syndrome. The smaller *~220kb* deletion (*β* = 4.5 kg/m^2^ [95%CI: 2.7-6.3 kg/m^2^], *p* = 1.8×10^−6^, AC=35) has been associated with early onset obesity, and spans *ATXN2L*, *TUFM*, *SH2B1*, *ATP2A1*, *RABEP2*, *CD19*, *NFATC2IP*, *SPNS1*, and *LAT*, with *SH2B1* the suspected causal obesity gene^3^. Obesity is also a phenotypic consequence of a larger *~593kb* deletion (β = 7.8 kg/m^2^ [95%CI: 6.2-9.4 kg/m^2^], *p* = 5.0×10^−23^, AC=58), which is further associated with neurodevelopmental delay and related conditions^4^. However, this deletion spans a wholly distinct set of genes which are suspected to play complex dosage-dependent roles in the phenotypic consequences of the syndrome^40^. As both subtypes of *16p11.2* deletion syndrome may present in early childhood, it is noteworthy that the effect we measure on BMI is in a cohort comprised entirely of older individuals, indicating burden of adult disease associated with the CNV locus.

After controlling for multiple comparisons, burden testing for BMI identifies a group of genes at chromosome *22q11.2* and recapitulates the list of genes affected by each of the *16p11.2* deletions. Variation at the *22q11.2* locus also constitutes one of the first named microdeletion syndromes, DiGeorge syndrome, which has variable phenotypic consequences including craniofacial dysmorphisms and conotruncal congenital heart disease, along with increased risk for an adverse cardiovascular outcomes and neuropsychiatric disease later in life^5^. Among individuals affected with *22q11.2* deletion syndrome obesity is a recognized manifestation of disease^42^, and we estimate a 0.3-0.4 point increase in BMI for genic CNVs near *22q11.2*. as well as 0.55-0.7 for genic CNV at *16p11.2*. The presence of these associations in a large volunteer cohort offers further evidence that syndromic alleles may contribute to the risk of common diseases in the general population.

Phenome-wide associations for each of the CNVs at *16p11.2* further highlight changes in biomarkers, biomeasures, and increased risk of common disease, consistent with high BMI over the course of a lifetime (Figure 4). Genome-wide significant phenotypes for the 220*kb* CNV recapitulate the established syndromic effects from early onset obesity. We observe significant increases, on the order of one standard deviation, in weight, BMI, hip and waist circumference, reticulocyte count, and comparative body size at age 10 for these individuals. The larger 593*kb* CNV associates with similar measures of body size and fat, as well as hypertension, diabetes, and abdominal hernia. These results are also indicative of effects due to developmental delay; namely, decreased measures of memory, higher Townsend deprivation, and lower lung capacity (FEV, FVC) with higher associated risk of respiratory failure. Taken together these results highlight the variable expressivity of CNV-related disease, as well as its long-term effects across the medical phenome.

**Figure 2:**
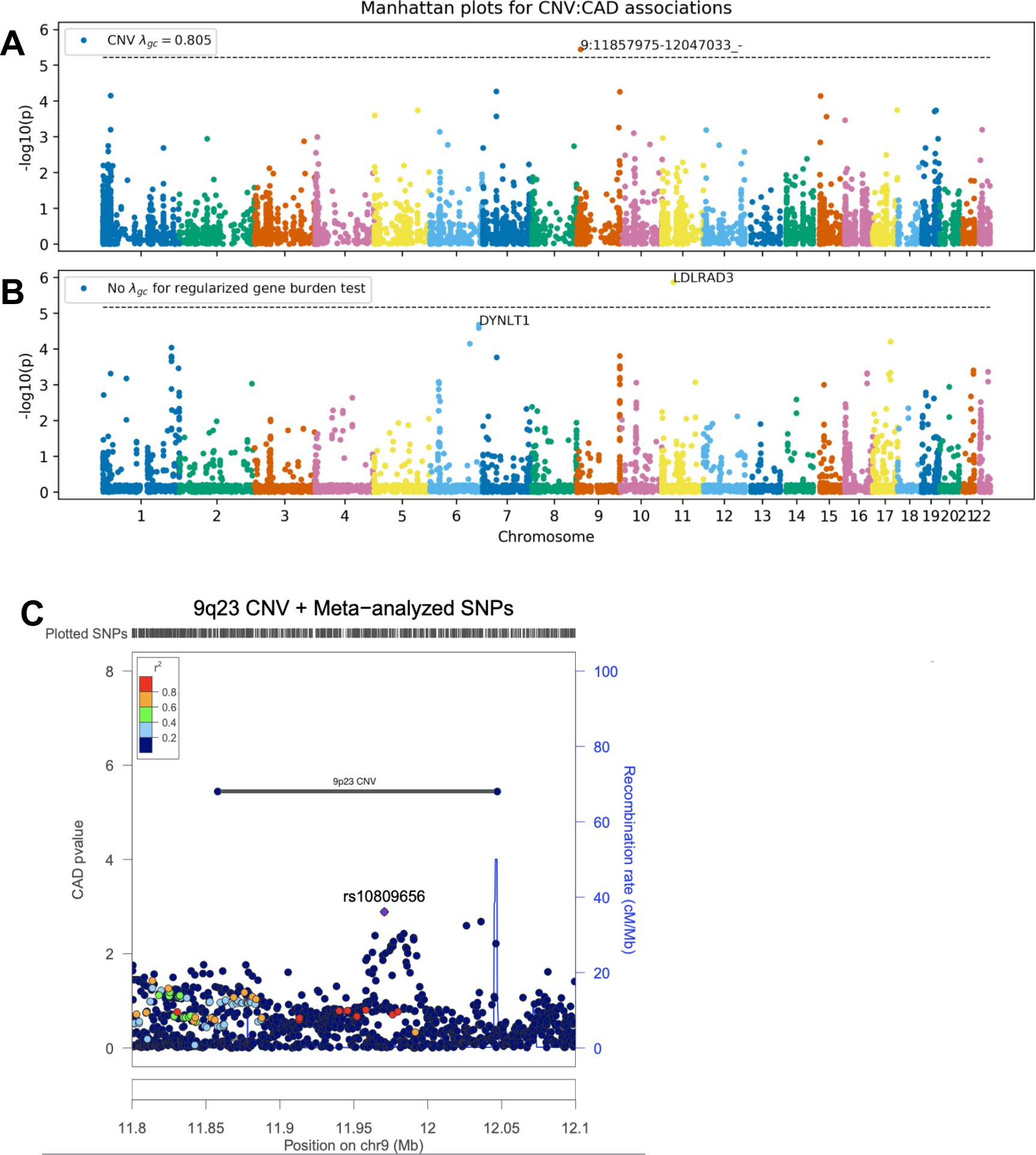
Genome-wide CNV associations for acute coronary artery disease (CAD). Manhattan plots for **(A)** genome-wide association of common copy number variants, and **(B)** genome-wide burden test of rare variants. **(C)** LocusZoom^38^ of 9p23 CNV and summary statistics from meta-analysis of CAD^33^ colored by marker LD with lead regional GWAS SNP (rs10809656) in HapMap^39^ European samples.

**Figure 3:**
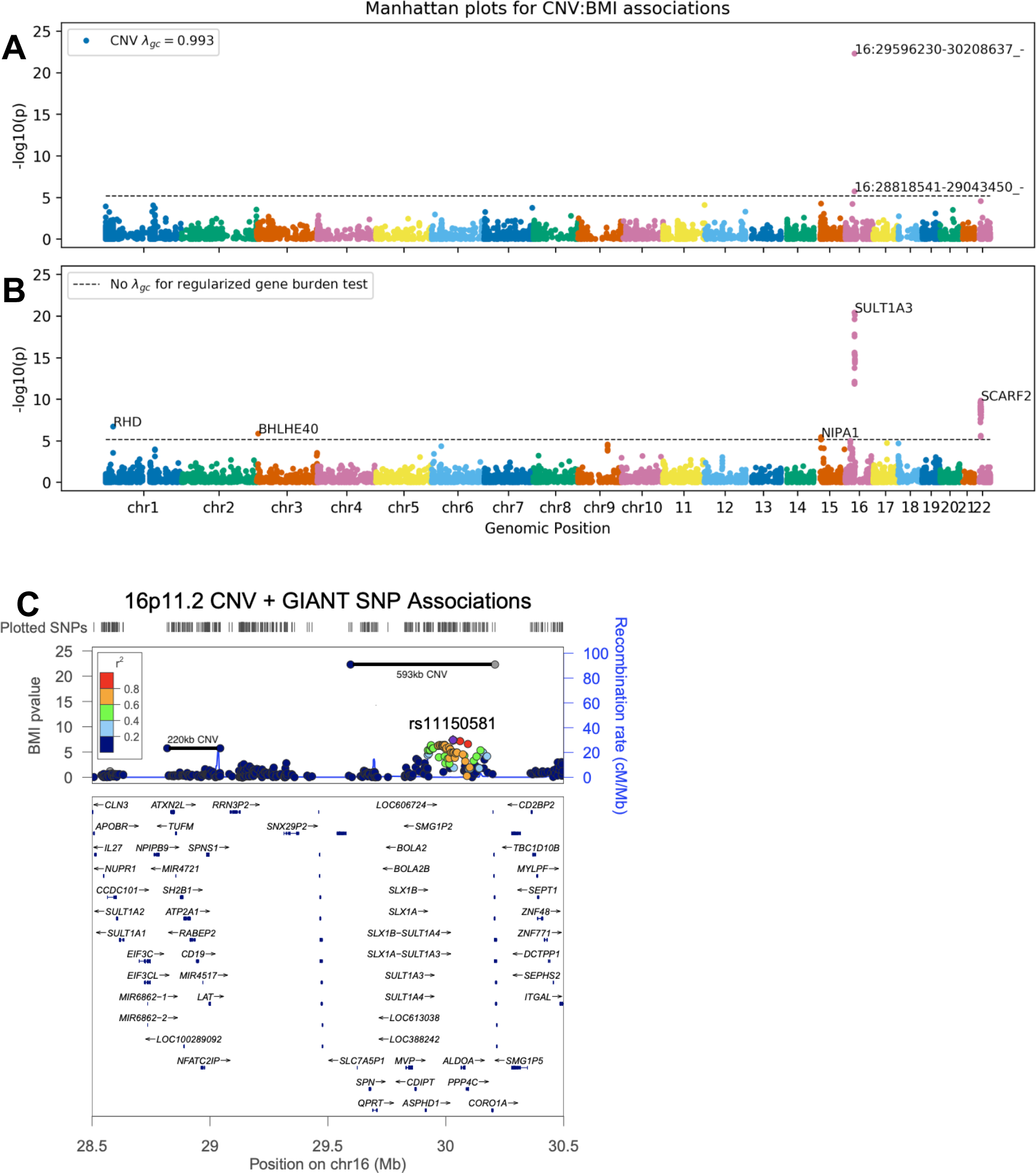
Genome-wide CNV associations for body mass index (BMI). Manhattan plots for **(A)** genome-wide association of common copy number variants, and **(B)** genome-wide burden test of rare variants. **(C)** LocusZoom of *16p11.2* CNVs and BMI summary statistics from the GIANT study^41^, colored by marker LD with the lead SNP at the locus (rs11150581), computed from HapMap European samples.

**Figure 4:**
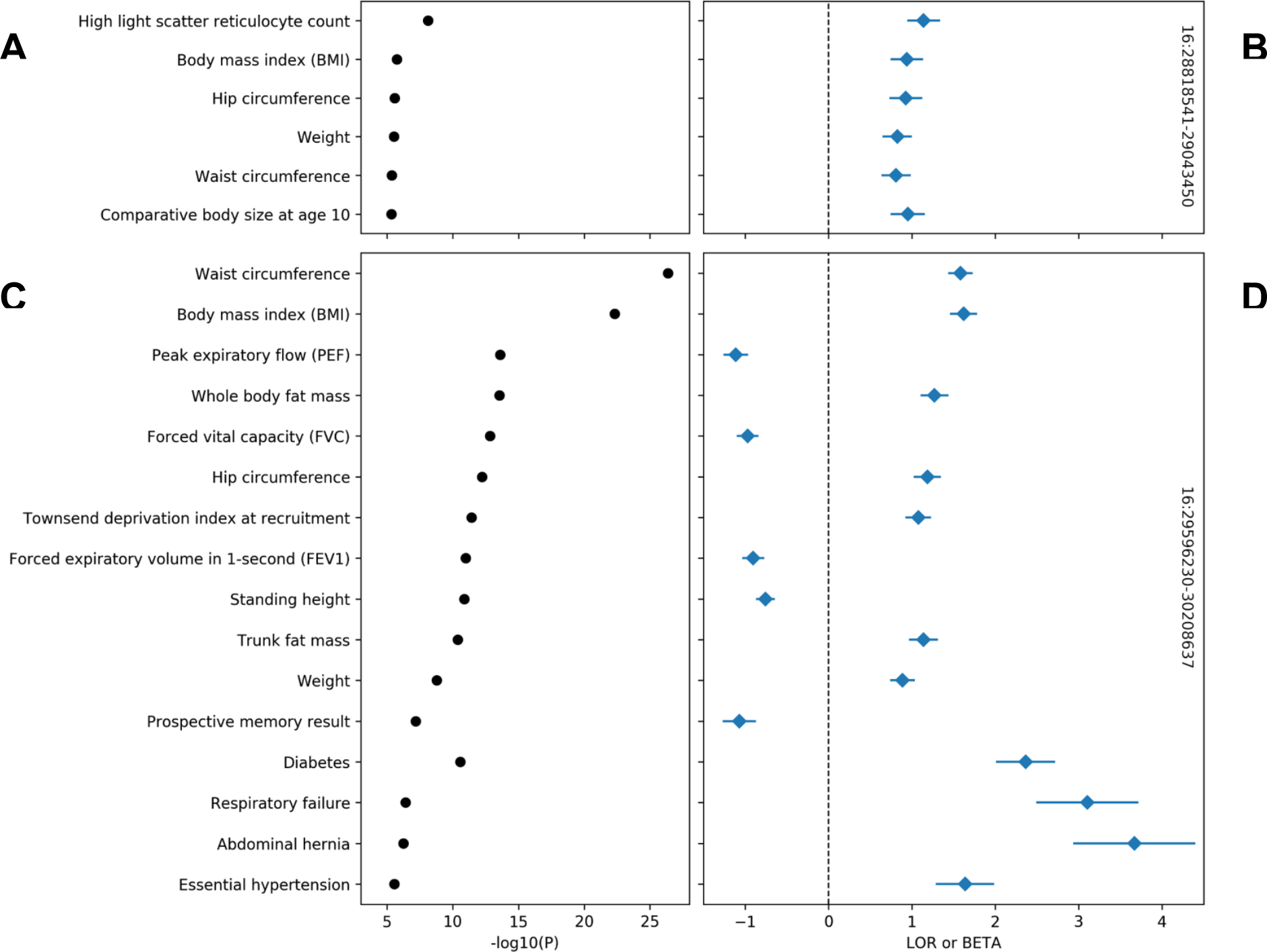
PheWAS of 16p11.2 CNVs. Selected genome-wide significant (*p* < 6×10^−6^) associations for 220*kb* (top panels) and 593*kb* (bottom panels) *16p11.2* CNVs. Traits are grouped by type (binary/quantitative) then sorted by *p*-value (left panels). Log-odds ratio and standardized betas (right panels) align with trait names on the y-axis, with the horizontal dashed line separating positive and negative direction of association.

## Discussion

In calling copy number variants and performing genetic association studies at scale from a large cohort of array-genotyped individuals with richly annotated phenotype data, we provide a portrait of the phenome-wide burden of genomic copy number variation. Our estimates of the individual-level burden of CNV and population-wide allele frequencies are consistent with previous reports, the deep phenotypic information available in the UK Biobank permits more finely tuned measures of the genic intolerance to CNV which include estimates of variation absent from our cohort of predominantly healthy, middle-aged individuals. We consider both rare and common CNVs in our association studies and identify effects of previously known (*22q11.2, 16p11.2*) syndromes and potentially novel (*9p23*) loci which have not yet been characterized or associated with disease.

Our study has significant limitations which inform our analysis. While arrays are an inexpensive way to genotype large cohorts, permitting straightforward algorithms to infer the presence of structural variation, the resulting CNV calls are limited by the density and placement of markers across chromosomes. For UK Biobank genotyping arrays in particular, there are large portions of genomic sequence with low marker density (in particular near centromeric regions) which bias our resulting genotype calls away from such regions. Array-derived CNV calls also suffer from limitations, in their inability to differentiate other structural events like inversions or translocations, or to determine breakpoint position at base-pair resolution. Complicating these barriers is the fact that our sample was genotyped on two distinct arrays, which may cause identical CNVs to present with different breakpoints across individuals in the call set. Our choice to present gene-level burden tests which include the vast majority of variants included in our CNV GWAS was informed by this realization.

Our associations are also heavily impacted by a known “healthy-cohort” bias, which results in null results for several phenotypes with known genetic contributions; notably, there are no hits in our burden tests for cancers other than leukemia and lymphomas. With this in mind, our constraint scores constitute a sobering observation of genetic survivorship bias. Estimates of gene-level intolerance to structural variation are derived from people who did not enroll in UK Biobank; the absence of individuals who were not healthy enough to participate or did not survive until age 40 constitutes a enrollment bias against severe early-onset disease. We take this opportunity to honor these non-participating individuals and their implicit contribution to our understanding of genetic disease. The observation of selection bias colors the interpretation of genetic findings from UK Biobank in general, as the cohort is relatively depleted of disease of early-onset morbidity and mortality and any genetic variation associated with these diseases will likewise be difficult to detect. While UK Biobank is unprecedented in size, scope, and scientific yield, our data illustrate that the anticipated findings from the proliferation of large biobank studies around the world will be influenced by implicit and explicit barriers to participation.

Despite selection against high-penetrance alleles causing early-onset disease, we detect a novel and strong association for coronary artery disease at *LDLRAD3*. While this locus has prior putative association with bone mineral density^43^, existing large-scale GWAS do not detect a strong association with coronary artery disease or established cardiometabolic risk-factors. However, the absence of gene level intolerance to truncating and missense variation in *LDLRAD3* does not suggest a compelling rationale for important biological function or role in disease. In our study, CNVs at this locus are associated with some established cardiometabolic risk factors, such as diabetes onset, smoking status, and arterial stiffness, but not obesity or other fat-related phenotypes (Figure S6). Consistent with our findings that a decrease in *LDLRAD3* dosage increases the risk of disease, a strong eQTL increasing *LDLRAD3* expression decreases the risk of disease when used as an instrument in a two-sample mendelian randomization in a large-scale study of coronary artery disease. Thus, our findings of an mRNA dosage effect are replicated at the gene level. These results highlight the utility of analyzing genic CNV which, when directly impacting mRNA dosage, offer a more easily interpretable mechanism distinct from alterations of protein structure or small changes in transcriptional regulation.

The observation of variation at the *16p11.2* and *22q11.2* loci sheds further light on the penetrance of syndromic loci in the general population. The *16p11.2* recurrent microdeletion syndrome was first described in individuals with autism and neuropsychiatric disease and may include seizures, brain and other anatomic abnormalities. Even accounting for the enrollment bias inherent to the UK Biobank our PheWAS detects a modest relationship to neurocognitive measurements via secondary markers of intellectual differences such as prospective memory for one variant at this locus. People carrying variation at the 22q11.2 locus within the general population are known to be at increased risk of neuropsychiatric diseases^44^ for which variable phenotypic penetrance is well recognized^345^. To wit, individuals with genetic variation at both syndromic loci were by and large sufficiently healthy and capable of volunteering to participate in the Biobank. Our findings support a growing recognition that the penetrance and effect sizes of syndromic alleles will likely require revision in the context of broad population-based surveys of rare genetic variation^46,47^.

Our findings add to the growing body of literature measuring global burden of structural variation across healthy and diseased individuals. Our estimates of the effects of common CNV suggest a notable role of structural variation in population-wide burden of common disease, and suggest genomic loci where novel CNV-derived syndromic disease may exist. For rare variants, our data offer broad phenotypic characterization of the effects of gene-specific knockouts, which may inform development of pharmacological and genetic therapies. While the functional consequence and pathogenicity of missense, synonymous, and noncoding single nucleotide variation within a gene may be difficult to classify, the mechanism of most genic CNV are clear: a dosage effect upon mRNA transcription. This population-scale catalog of variation and the described associations with a multiplicity of diseases should be of immediate use by genetic clinicians in classification of novel and rare CNV detected in clinical testing. Full support for gene- and variant-level browsing is forthcoming in a future version of the Global Biobank Engine (biobankengine.stanford.edu). In the interim, summary statistics from association studies described here, as well as for all phenotypes present on the engine, are freely available for download on the site. We hope that these data will be leveraged to empower future analyses of the phenome-wide effects of structural variation and gene-level dosage effects.

## Acknowledgements

This research has been conducted using the UK Biobank Resource under application numbers 24983, 16698, 13721, and 15860. We thank all the participants in the study. The primary and processed data used to generate the analyses presented here are available in the UK Biobank access management system (https://amsportal.ukbiobank.ac.uk/) for application 24983, “Generating effective therapeutic hypotheses from genomic and hospital linkage data” (http://www.ukbiobank.ac.uk/wp-content/uploads/2017/06/24983-Dr-Manuel-Rivas.pdf), and the results are displayed in the Global Biobank Engine (https://biobankengine.stanford.edu).

M.A.R. is supported by Stanford University and a National Institute of Health center for Multi- and Trans-ethnic Mapping of Mendelian and Complex Diseases grant (5U01 HG009080). This work was supported by National Human Genome Research Institute (NHGRI) of the National Institutes of Health (NIH) under awards R01HG010140. The content is solely the responsibility of the authors and does not necessarily represent the official views of the National Institutes of Health.

## Methods

CNVs were called using PennCNV v1.0.4 on raw signal intensity data from each genotyping array. Phenotype data was derived from data-fields collected for UK Biobank corresponding to various body measurements, biomarkers, disease diagnoses and medical procedures from medical records, as well as a questionnaire about lifestyle and medical history. Summary-level data from all statistical tests described here, as well as more thorough documentation on phenotyping, will be available on the Global Biobank Engine^14^ (biobankengine.stanford.edu) and can be found in related publications^48^.

### CNV calling in UK Biobank

Methods for genetic data acquisition and quality control as performed by the UK Biobank have been previously described^13^. In brief, two similar arrays were used for targeted genotyping within the study population: the UK BiLEVE Axiom Array (*n*=49,950) by Affymetrix and the UK Biobank Axiom Array (*n*=438,427), which was custom-designed by Applied Biosystems. Samples and array markers were subject to threshold-based filtration and quality control prior to public release. Specifically, markers were tested for discordance across control replicates, departures from Hardy-Weinberg equilibrium, as well as effects due to batch, plate, array, and sex; affected markers were set as missing in affected batches or removed. Similarly, samples were tested for missingness (>5%) and heterozygosity across a set of high-quality markers, but samples identified as low quality (*n*=968) were not excluded. We also chose to include these samples in this analysis, considering that large structural variants may have been responsible for their poor quality with respect to metrics used for filtration.

We used PennCNV *v1.0.4*^15^ to call CNVs within each of the 106 genotyping batches from UK Biobank. We first estimate genomic runs of heterozygosity (RoH) for each sample using a previously developed pipeline in PLINK^49,50^ using the --homozyg option. We then select *n*=100 samples with total RoH covering less than 20MB to train a hidden markov model (HMM) of copy state on each chromosome. HMM training was initialized with conditions optimized for Affymetrix arrays (affygw6.hmm), provided in PennCNV resources. We used the general calling mode, which performs likelihood-based testing for copy-number state (CN=0,1,2,3,4) at each input marker using its log-normalized signal intensity and allele balance in a given sample. We also apply adjustment for GC content across sites using waviness factor correction^51^. After CNV calling, we exclude 1,360 samples with over 30 called CNVs from downstream analysis, resulting in a cohort of 472,228 individuals with 278,455 unique variants.

### Gene-level constraint estimation

Regional selective constraint to CNV was estimated for all protein-coding genes, with genic CNV defined as any variant overlapping within 10*kb* of the HGNC gene region. We estimate a null model of structural mutation empirically, and model burden of genic CNV as a linear function of gene size, fraction of genic sequence covered by regions of segmental duplication as extracted from the UCSC Genome Browser^52,53^. We also account for biased observations due to array genotyping by including the number of genic markers as a covariate. The formula for this null model can be written as:

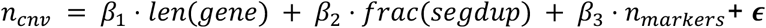

From this model, we compute constraint *z-*scores for each gene using its negated standardized residual for each gene, winsorizing the negative tail at the lowest 5% of values. We also compute the probability of loss of function intolerance (pLI) as the non-normalized residual over the number of expected CNV, with negative values rounded to zero.

### Genetic associations

Variant-level associations in UK Biobank were estimated with PLINK v2.00a (5 Jan 2018). We used the --glm firth-fallback option for computation. This option is a hybrid algorithm for logistic regression which defaults to a standard regression solver for computation, falling back to Firth’s regression (https://cran.r-project.org/web/packages/logistf/index.html) in cases where one of the cells of the 2×2 contingency table is zero, or where the traditional method fails to converge in a pre-specified number of iterations. These analyses were performed in a subset of 337,538 unrelated individuals of self-reported white British ancestry, and were controlled for age, sex, and 4 marker-based genomic principal components from the UK Biobank PCA calculation. To ensure adequate power for estimating genetic effects, we perform these tests on 8,274 CNVs observed at a frequency of 0.005% (1 in 20,000, or 18 individuals) in the whole sample of individuals with called CNVs.

Gene-level burden tests were conducted across all gene:phenotype pairs for genes with at least 5 individuals with overlapping CNV. Genic burden was encoded as a binary variable which indicates whether an individual has a CNV which contains any overlap within 10,000 base pairs of the HGNC gene region. CNVs which overlapped several gene regions were used for analysis in each gene. We treat deletions and duplications identically, with the assumption that any CNV which overlaps a gene in this fashion will disrupt its normal function. Effects of genic CNV burden were estimated by linear and regularized logistic regression with the python package Statsmodels, respectively computed using the .fit() and .fit_regularized() methods. We included the following as covariates in both models: age, sex, four marker-based genomic principal components from UK Biobank’s PCA calculation, and the number and combined length of CNVs in each individual.

Two-sample mendelian randomization was performed via the MR Base web app using GWAS summary statistics for *LDLRAD3* expression QTLs from a CARDIoGRAMplusC4D meta-analysis^37^. We report Wald summary statistics from inverse-variance weighted Egger regression; these are the default analysis options for the web interface.

**Figure S1:**
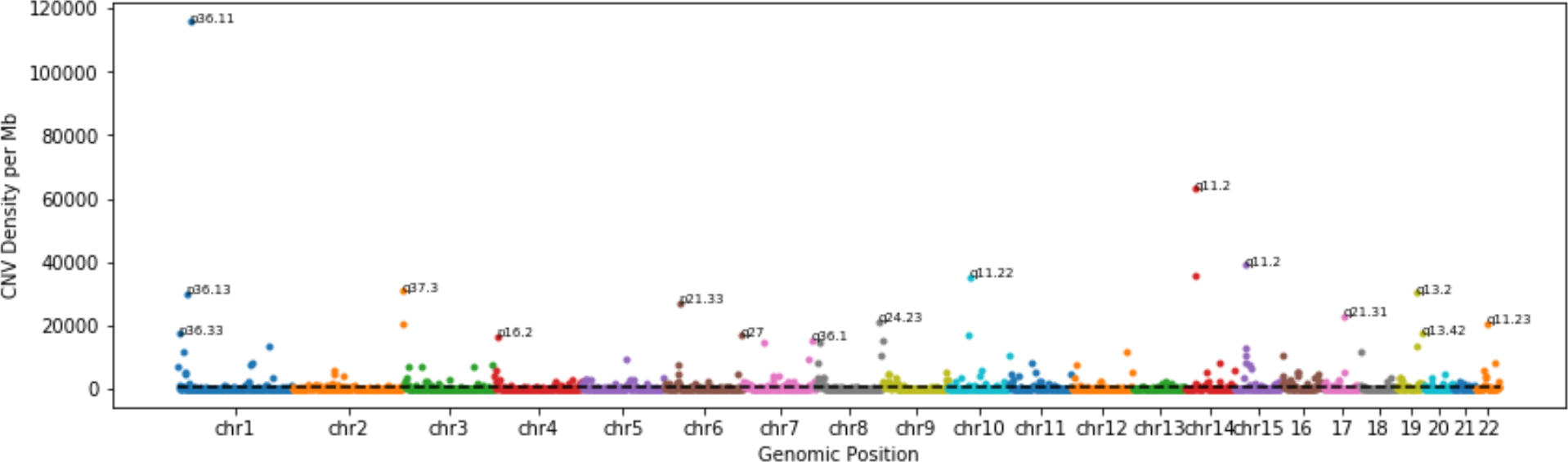
CNV density weighted by allele count in UK Biobank. Per-megabase genomic density of CNV, weighted by number of observations across all samples in UK Biobank. Variants are counted by whether the CNV has any overlap with 10 megabase (Mb) windows tiling each chromosome. Selected hotspots of structural variation are labeled by the region’s corresponding cytogenic band.

**Figure S2:**
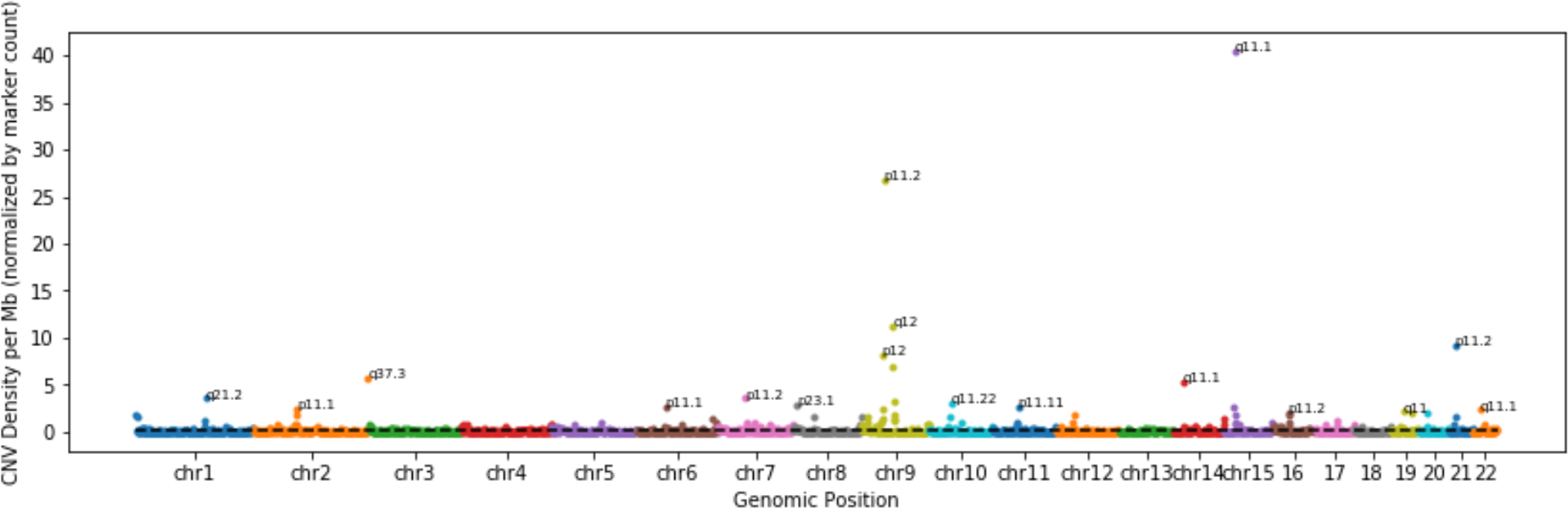
CNV density normalized by array marker density in UK Biobank. Variants are counted by whether the CNV has any overlap with 10 megabase (Mb) windows tiling each chromosome, then divided by the number of markers in the window. Regions with no array markers are defined to have density of zero. Selected hotspots of structural variation are labeled by the region’s corresponding cytogenic band.

**Figure S3:**
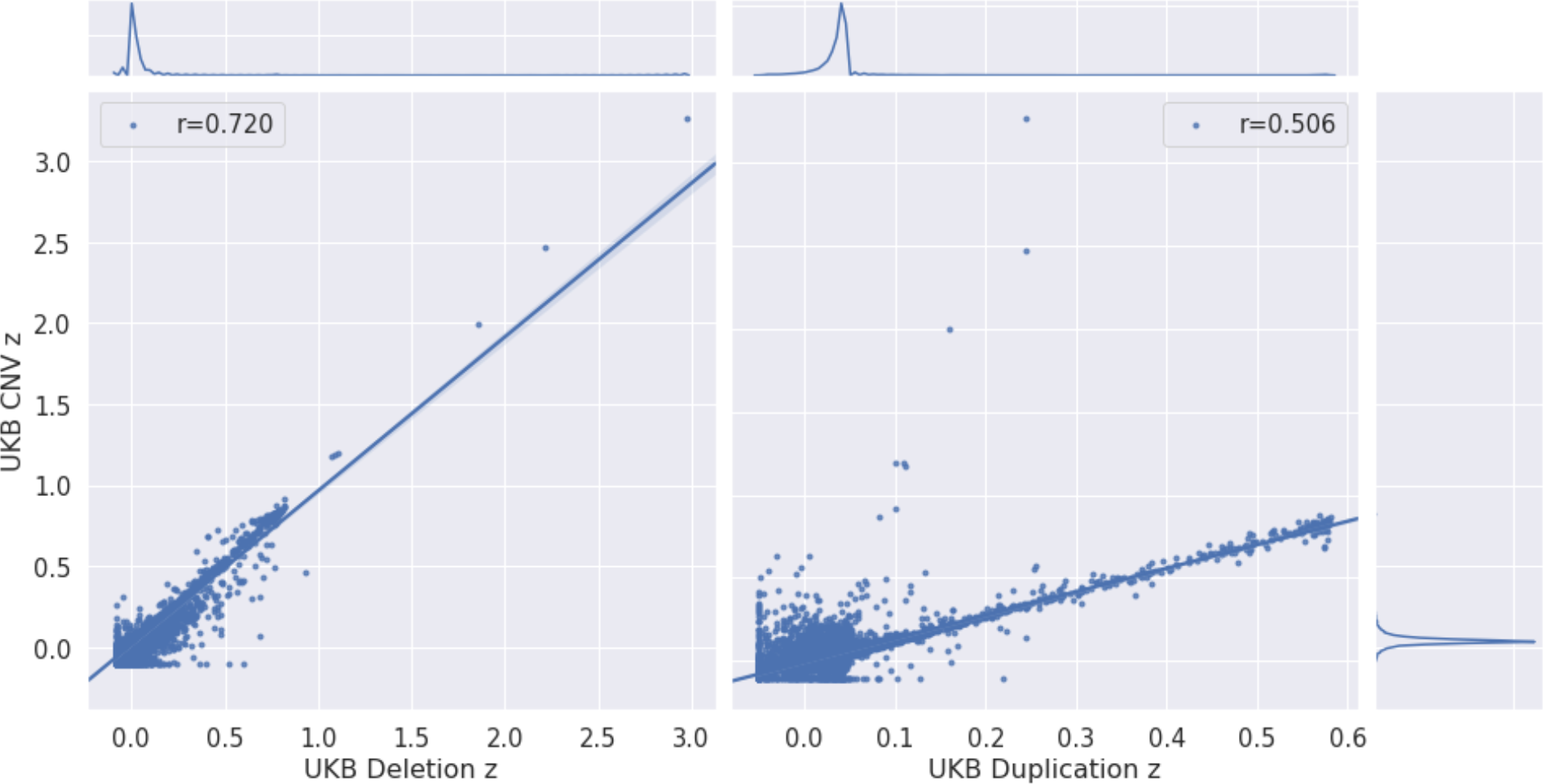
Correlation between intolerance measures for partial-gene deletion, whole-gene duplication, and CNV burden. The legend for each panel denotes correlation (Spearman’s *r*) between burden-constraint and each other measure. Kernel density estimates for each distribution of constraint scores are in the panels opposite their corresponding axis labels.

**Figure S4:**
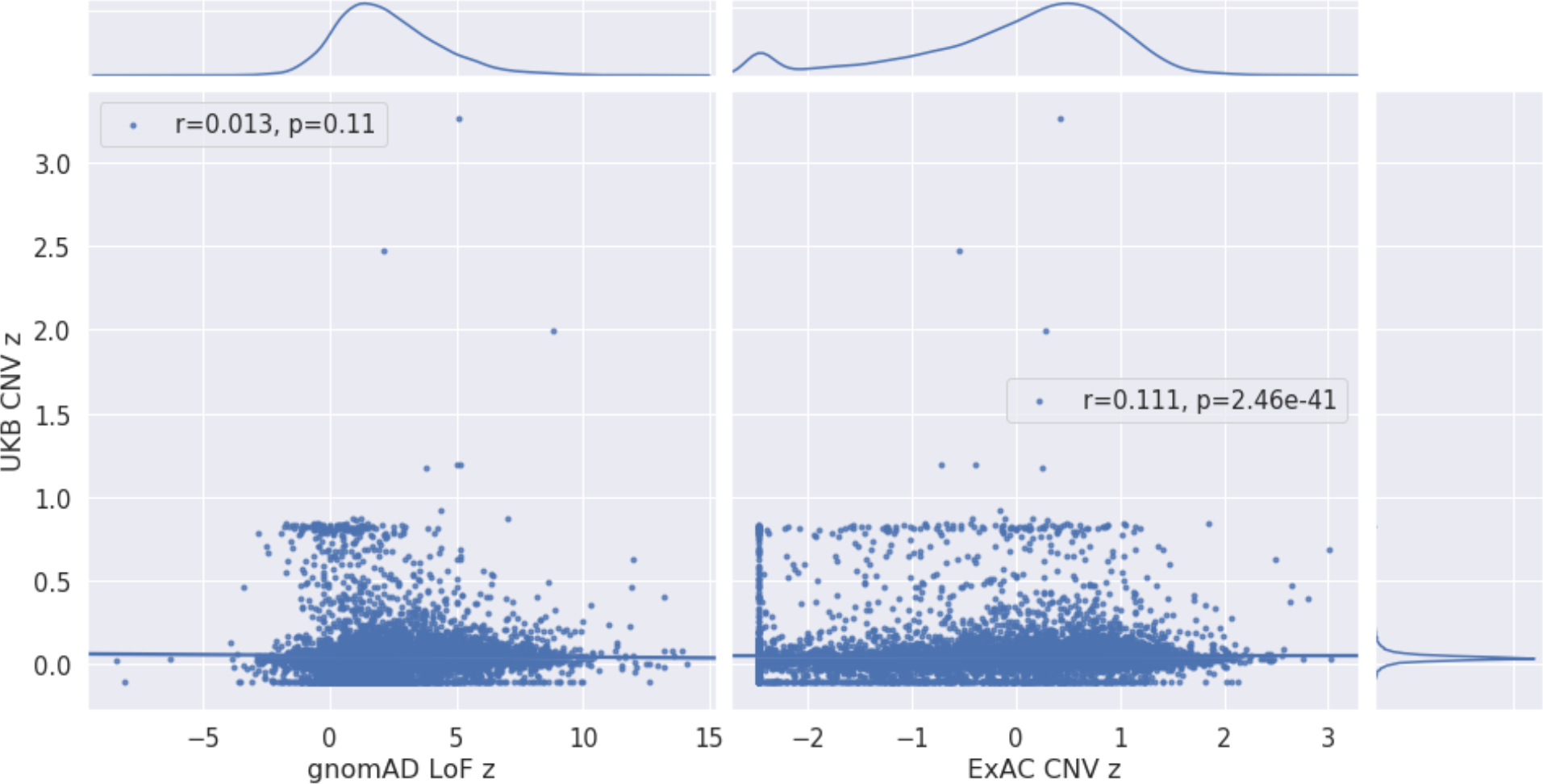
Distribution of constraint z-scores from UK Biobank and ExAC/gnomAD. Our measures of gene-level intolerance to structural variation show nominal correlation with gnomAD loss of function constraint *z*-scores (Spearman’s *r* = 0.013, left), and modest correlation with CNV-intolerance in ExAC (Spearman’s *r* = 0.11, right panel). Gaussian kernel density estimates for each distribution of *z*-scores are opposite their corresponding axes. While correlation between constraint measures across datasets is non-random, we suspect cohort-specific effects and varying definitions of genic burden of variation drive these departures. As a cohort of predominantly healthy adults, intolerance to variation in UK Biobank constraint is driven by severe early onset disease, while the same measures in ExAC/gnomAD, whose samples have a more diverse age range and relatively higher of burden of disease, highlight genes involved with fundamental biological processes whose loss of function likely confer phenotypic consequences causing embryonic lethality.

**Figure S5:**
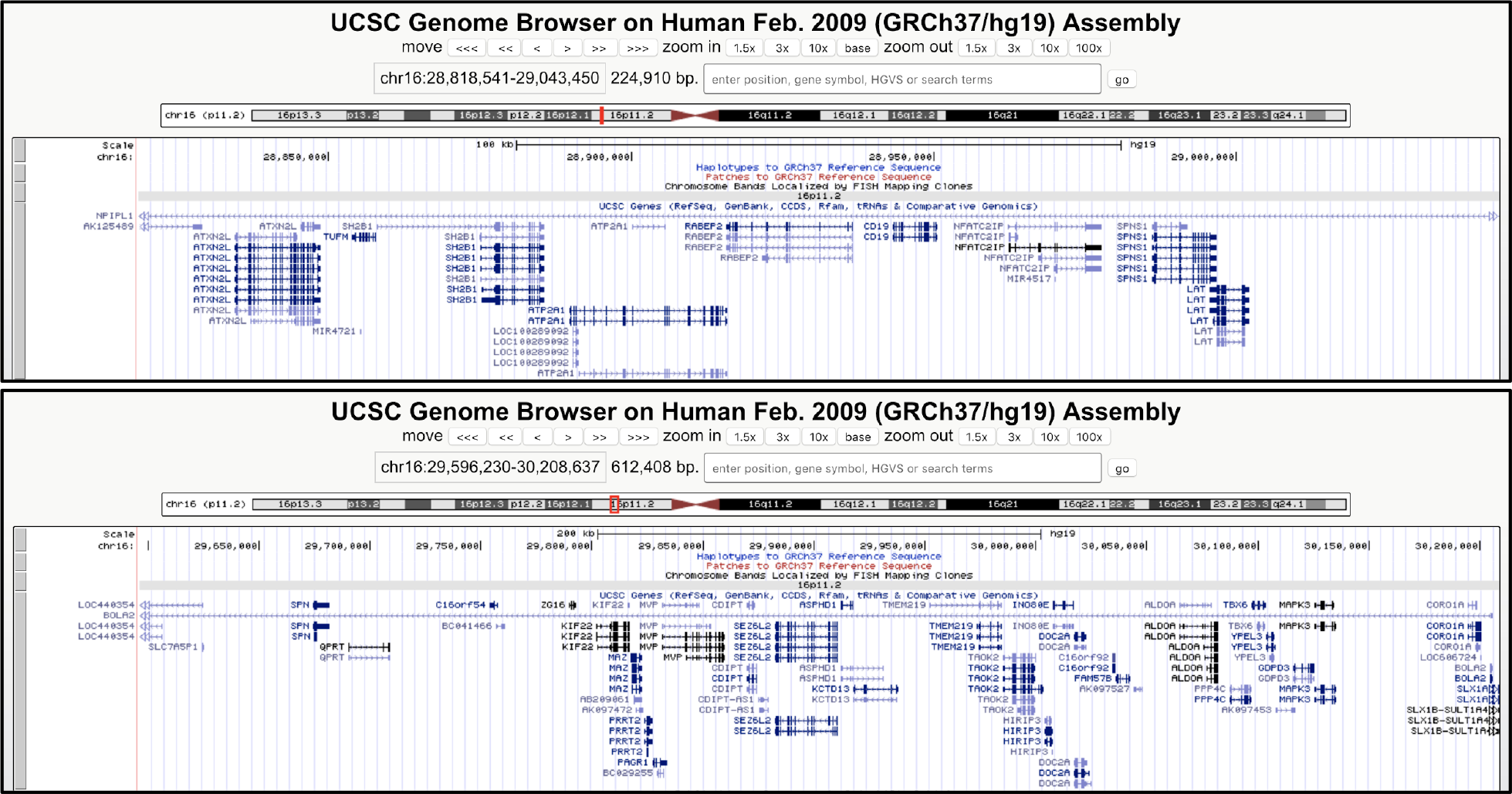
UCSC Genome Browser tracks for *220kb* (top panel) and *593kb* (bottom panel) CNVs at *Chr16q11.2*.

**Figure S6:**
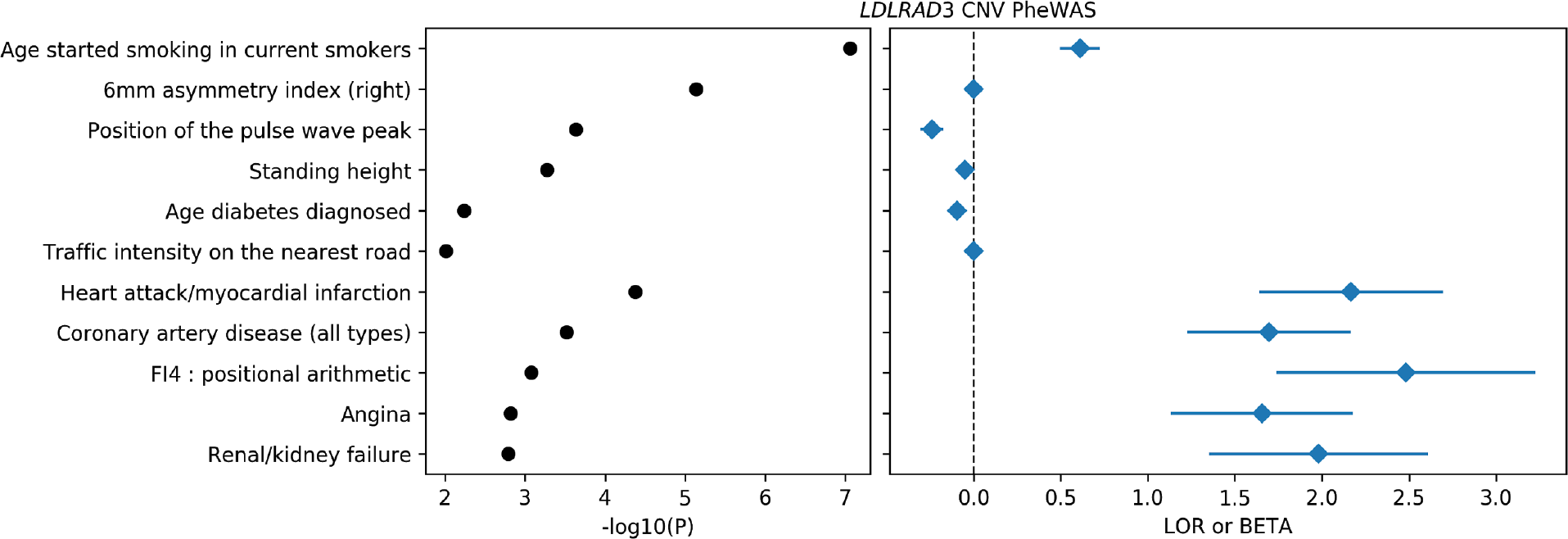
*LDLRAD3*burden test PheWas. Significant (*p* < 10^−3^) associations between regularized burden tests for *LDLRAD3* CNV and phenotypes. We highlight quantitative traits with *n* > 15,000 observations and binary traits with *n* > 100 cases. Traits are grouped by data type then sorted by *p*-value (left). Log-odds ratio and standardized betas (right; for binary and quantitative traits, respectively) align with trait names on the y-axis, with the horizontal dashed line separating positive and negative direction of association.

**Table S1:** 15 genes most intolerant to overlapping deletion (left), and whole-gene duplication (right), with respective constraint z-scores.

|  | Deletion z |
| --- | --- |
| BRCA2 | 2.974 |
| BRCA1 | 2.212 |
| APC | 1.857 |
| ATM | 1.104 |
| MSH2 | 1.087 |
| MLH1 | 1.072 |
| MYH7 | 0.934 |
| PMS2 | 0.880 |
| TTN | 0.833 |
| PVRL2 | 0.823 |
| MSH6 | 0.821 |
| SBDS | 0.816 |
| CYP3A4 | 0.816 |
| SPATA31D1 | 0.815 |
| OTOP1 | 0.809 |

|  | Duplication z |
| --- | --- |
| HLA-DRB1 | 0.581 |
| FRG2B | 0.581 |
| SPATA31D1 | 0.581 |
| SLC35G6 | 0.580 |
| NAT8 | 0.580 |
| TUBB8 | 0.580 |
| CSH2 | 0.580 |
| ZNF302 | 0.580 |
| CSHL1 | 0.580 |
| GH1 | 0.580 |
| CGB2 | 0.579 |
| FAM215A | 0.579 |
| OR4F17 | 0.579 |
| CGB5 | 0.579 |
| CGB7 | 0.579 |

**Table S2:** GO pathways most intolerant to overlapping deletion (top), and whole-gene duplication (bottom), with change in constraint z-scores and significance thereof (t-test) relative to other pathways.

| | Deletion-intolerant Pathway | $\Delta z$ | P |
| --- | --- | --- | --- |
| GO:0045095 | keratin filament | 0.23 | $2.22 \times 10^{-28}$ |
| GO:0000137 | Golgi cis cisterna | 0.36 | $2.10 \times 10^{-26}$ |
| GO:0052697 | xenobiotic glucuronidation | 0.44 | $5.29 \times 10^{-20}$ |
| GO:0008194 | UDP-glycosyltransferase activity | 0.32 | $4.79 \times 10^{-19}$ |
| GO:0005515 | protein binding | 0.05 | $1.19 \times 10^{-17}$ |
| GO:0000800 | lateral element | 0.39 | $1.36 \times 10^{-17}$ |
| GO:0031424 | keratinization | 0.15 | $1.33 \times 10^{-16}$ |
| GO:0015020 | glucuronosyltransferase activity | 0.31 | $2.56 \times 10^{-16}$ |
| GO:0042954 | lipoprotein transporter activity | 0.37 | $3.33 \times 10^{-14}$ |
| GO:0005131 | growth hormone receptor binding | 0.50 | $3.48 \times 10^{-13}$ |
| GO:0008202 | steroid metabolic process | 0.24 | $4.49 \times 10^{-13}$ |
| GO:0046703 | natural killer cell lectin-like receptor binding | 0.39 | $7.08 \times 10^{-13}$ |
| GO:0008274 | gamma-tubulin ring complex | 0.32 | $1.08 \times 10^{-12}$ |
| GO:0005132 | type I interferon receptor binding | 0.28 | $1.40 \times 10^{-12}$ |
| GO:0052696 | flavonoid glucuronidation | 0.40 | $1.59 \times 10^{-12}$ |

| | Duplication-intolerant Pathway | $\Delta z$ | P |
| --- | --- | --- | --- |
| GO:0000137 | Golgi cis cisterna | 0.34 | $2.57 \times 10^{-46}$ |
| GO:0045095 | keratin filament | 0.19 | $4.45 \times 10^{-34}$ |
| GO:0005515 | protein binding | 0.05 | $1.56 \times 10^{-30}$ |
| GO:0031424 | keratinization | 0.14 | $2.12 \times 10^{-23}$ |
| GO:0008202 | steroid metabolic process | 0.22 | $1.60 \times 10^{-22}$ |
| GO:0005132 | type I interferon receptor binding | 0.27 | $8.04 \times 10^{-20}$ |
| GO:0008194 | UDP-glycosyltransferase activity | 0.24 | $5.47 \times 10^{-19}$ |
| GO:0005801 | cis-Golgi network | 0.17 | $6.16 \times 10^{-19}$ |
| GO:0046703 | natural killer cell lectin-like receptor binding | 0.37 | $6.81 \times 10^{-19}$ |
| GO:0052697 | xenobiotic glucuronidation | 0.30 | $9.68 \times 10^{-18}$ |
| GO:0002323 | natural killer cell activation involved in immune response | 0.27 | $1.49 \times 10^{-17}$ |
| GO:0033141 | positive regulation of peptidyl-serine phosphorylation of STAT protein | 0.24 | $7.73 \times 10^{-17}$ |
| GO:0006805 | xenobiotic metabolic process | 0.16 | $1.73 \times 10^{-15}$ |
| GO:0005131 | growth hormone receptor binding | 0.39 | $2.39 \times 10^{-15}$ |
| GO:0042271 | susceptibility to natural killer cell mediated cytotoxicity | 0.24 | $4.69 \times 10^{-15}$ |

